# pyMultiVideo: Open-source, Python-based, software for acquiring, synchronising, and compressing video from multiple cameras

**DOI:** 10.64898/2026.11.23.768693

**Authors:** Alif Aziz, Thomas Akam

## Abstract

Video acquisition plays a critical role in many scientific experiments, enabling the quantification of human or animal behaviour, and movement of physical systems. Scientific applications place exacting demands on video acquisition, often requiring multiple cameras, high frame rates, synchronisation between cameras or with other hardware, control of camera parameters, and efficient data storage with comprehensive metadata. Here we present pyMultiVideo, an open-source Python application for scientific video acquisition. Features of the software include a graphical user interface (GUI) for efficiently configuring and running multi-camera experiments, online video compression using the computer’s GPU, and synchronisation through recording GPIO pins states or external frame triggering. The code-base is simple and modular, with a separation of camera API from GUI code to facilitate adding support for additional camera systems. We outline the design and functionality of the software and present validation experiments demonstrating simultaneous recording and compression from multi-camera setups, at high frame rates without dropped frames on modest computer hardware.

## Introduction

Video acquisition is an key tool across many scientific disciplines, including the study of human and animal behaviour. The development of deep-learning based tools for pose estimation and key-point tracking ^1–3^, and for identifying behavioural states from video data ^4–6^, has dramatically increasing the power of video for quantifying behaviour, and made video analysis ubiquitous in contemporary behavioural sciences and neuroscience.

Scientific video acquisition necessitates a demanding set of software features. Experiments often use multiple cameras, for 3D tracking or to record multiple subjects, and may require high frame rates for precise temporal resolution. This generates large data which require compression for efficient storage, ideally performed online during acquisition to avoid the need for specialised storage hardware and post-processing. For reproducibility and consistency, camera acquisition parameters such as exposure and gain must be controlled and recorded as metadata. As video is often one of multiple data streams generated by an experiment, mechanisms are needed for synchronisation, both between cameras and with other hardware systems. The software’s user interface is also important, both for ease of use and to minimise human error in configuring and running experiments.

Commercial software for multi-camera video acquisition exists, but often either lacks features needed for scientific applications or requires an expensive licence. A number of open source video recording applications have been developed for scientific use ^7–11^, with varying feature sets (see discussion), however there remains a need for user friendly and extensible software that supports the rich set of features required for scientific video acquisition.

To address this we developed pyMultiVideo; Python-based open source software that provides a solution for acquiring, visualising, synchronising and compressing video from multiple cameras. The userfriendly GUI implements functionality for saving and loading experimental configurations to facilitate high-throughput experiments while minimising the risk of user error, and detailed metadata is automatically saved with each video. We have made a substantial effort to ensure the code is well documented and structured, with a particular emphasis on making it straightforward to add support for new camera systems, and have created detailed online user documentation and a Github user discussion board. Camera API modules have been implemented for cameras from FLIR and Ximea, though the software has primarily been tested with FLIR Chameleon 3 cameras. Here we report the system design and rationale, as well as validation experiments characterising system performance.

## Results

### Graphical User Interface

The graphical user interface is implemented using PyQt6 and PyQtGraph ^12^, and organised into two tabs: *Video Capture* and *Settings*.

The *Video Capture* tab (Figure 1A) is used to control recording and display video feeds from multiple cameras. It has controls to select cameras, set subject IDs and the data directory, start and stop recording for individual or all cameras, configure the number of columns in the video feed layout, and for saving or loading the experiment configuration. Saving the experiment configuration saves the current state of the tab to a JSON file, which can subsequently be loaded to quickly and accurately configure acquisition for an experiment. Keyboard shortcuts are provided to maximise the video feeds within the window by hiding the controls, and to full-screen the video feeds so they fill the monitor. Information overlays on the video feeds display the current empirical frame rate, whether video is being recorded, and the current state of the GPIO pins, with warnings displayed if frames are being dropped.

**Figure 1:**
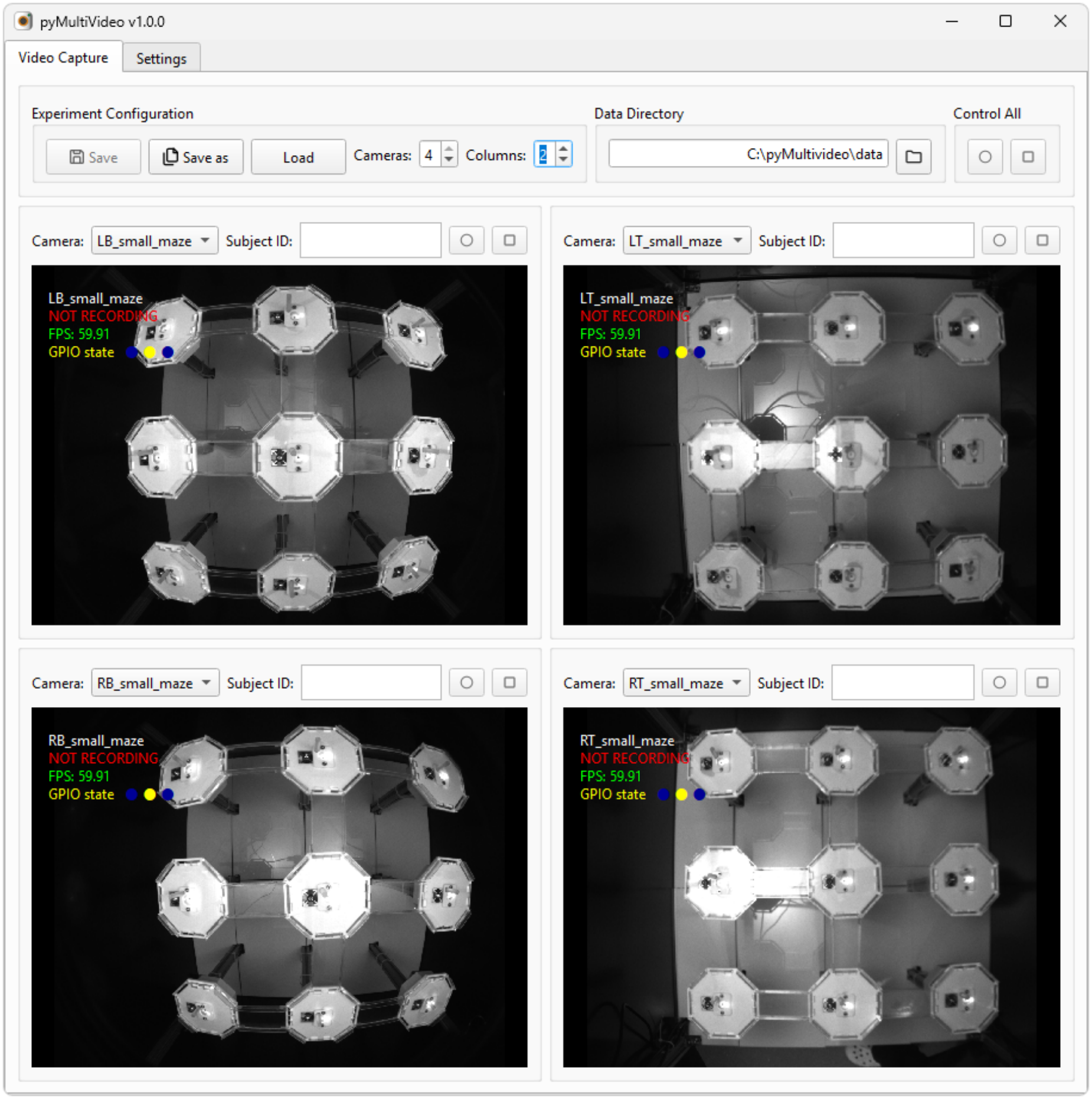
Video capture tab. used for configuring experiments and recording video.

The *Settings* tab (Figure 2) is used to configure camera and recording settings. For each camera, controls are provided to set the frame rate, exposure, gain, enable/disable external frame triggering, apply optional downsampling prior to saving video, and to specify a display name used to identify the camera in the GUI. The video feed from individual cameras can be previewed in the tab to visualise the effect of the current settings. Controls are also provided for specifying video compression parameters including the quality, encoding speed and compression standard. A further set of controls can optionally be used to configure pulse output from a microcontroller connected to the computer for use as a frame trigger signal (see Synchronisation below). Settings persist across closing and reopening the program using JSON files saved to disk.

**Figure 2:**
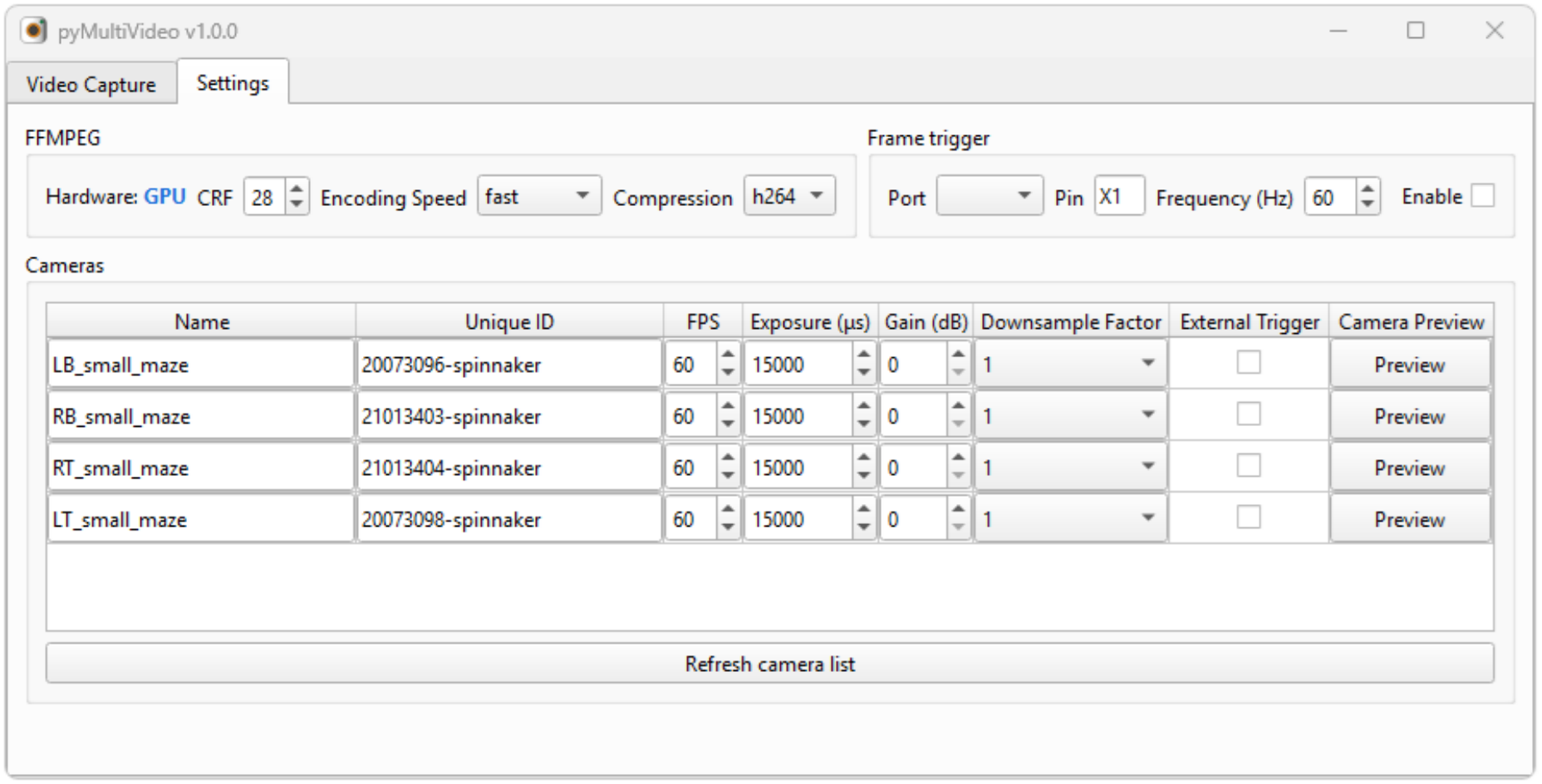
Settings tab. used for configuring camera and application settings.

This design implements a conceptual separation between experiment settings - i.e. the cameras used and their corresponding subject IDs and data directory, and acquisition settings - i.e. camera, frame triggering and video compression settings. The former are set in the video capture tab and manually saved to named experiment configuration files, while the latter are set in the settings tab and automatically persisted across sessions. The rationale for this is that in typical experimental workflows many different experiments are run on the same hardware setups using the same acquisition settings.

### Data Format

Each video recording creates three data files; *video data, metadata*, and *frame info*, with file names determined by the subject ID, recording start date-time, and file type.

Video data is saved as an .*mp4* file using h264 or h265 compression, selectable from the settings tab, with h264 used by default due to being most widely supported. This standard video file format can be straightforwardly loaded into most analysis tools and media players.

Video metadata is saved as a *JSON* file containing camera and acquisition settings, subject and camera IDs, recording start and end date-times, and the number of recorded and dropped frames. JSON is a lightweight data interchange format designed to be readable by humans and easy for computers to parse, that is supported by most programming languages.

The frame info file is a .*csv* with one row per recorded frame, with columns for the frame number, frame timestamp in microseconds relative to the first frame, and the state of the camera GPIO pins when the frame was acquired. Frame numbers and timestamps are extracted from the image metadata generated by the camera system and can be used to identify where any dropped frames occurred if present.

### Synchronisation

pyMultiVideo provides two primary mechanisms for synchronising video between cameras and with other hardware; external frame triggering, and reading camera GPIO pin states.

External frame triggering is enabled or disabled on a per-camera basis in the settings tab. When external triggering is disabled, frame acquisition is triggered internally by the camera using the frame rate set in the settings tab. When external frame triggering is enabled, the camera acquires a frame only when it receives a voltage pulse on its trigger input pin, such that the rate and timing of frame acquisition is determined by the timing of the trigger pulses.

pyMultiVideo implements functionality to output pulses from a microcontroller connected to the computer, which can be used to trigger frame acquisition by connecting the microcontroller pin used for pulse output to the frame trigger input pins on the cameras (ground pins must also be connected between micro- controller and cameras). The software automatically detects and configures connected microcontrollers running [MicroPython](https://micropython.org/), and the settings tab implements controls to select the serial port the microcontroller is on, specify the pin used for pulse output, the pulse frequency, and enable/disable pulse output. When recording is started or stopped on multiple cameras, trigger pulse output is paused as the software iterates over cameras to execute the command, ensuring that the same set of pulses trigger the recorded frames on all cameras.

Reading the state of camera GPIO pins provides an alternative mechanism for synchronising video with other hardware. Each time a frame of video is acquired from the camera, the corresponding GPIO pin states are read from the image metadata and saved to the frame info csv file. As the pin state is only recorded once each frame, the time resolution of the recorded signal is limited by the camera frame rate, so timing precision will be lower than that achieved by external frame triggering. The typical use case is to record sync pulses generated by other hardware systems, which can subsequently be used to align the video with other data streams. In our lab we output sync pulses from the pyControl ^13^ hardware that controls our behavioural setups, and route these to both the video camera and physiology recording systems. We use sync pulse trains with random inter-pulse-intervals which ensures a unique match between the inter-pulse-interval sequences recorded on each system, so it is always possible to identify which pulse corresponds to which across systems. For more information on this synchronisation approach, including a Python module for automatically aligning data using such sync pulses, see the pyControl [synchronisation](https://pycontrol.readthedocs.io/en/latest/user-guide/synchronisation/) docs.

A final possible approach to synchronisation is to use the recording start and end times recorded in the video metadata to convert frame timestamps into in the computer clock’s reference frame, to align video with other signals for which this time reference is available. This is unlikely to be as accurate as using sync pulses or external frame triggering, but may be sufficient when timing precision is not critical.

### Software Architecture

The pyMultiVideo codebase is organised into classes that implement different components of the GUI and camera control system. Figure 3 shows the organisation of the main classes involved in video acquisition. The Video Capture Tab instantiates one or more Camera Widgets each responsible for controlling and displaying video from a single camera. Each Camera Widget contains a Camera API instance which handles interactions with the camera hardware, and a Data Recorder which handles opening and closing data files and saving data to disk during recordings.

**Figure 3:**
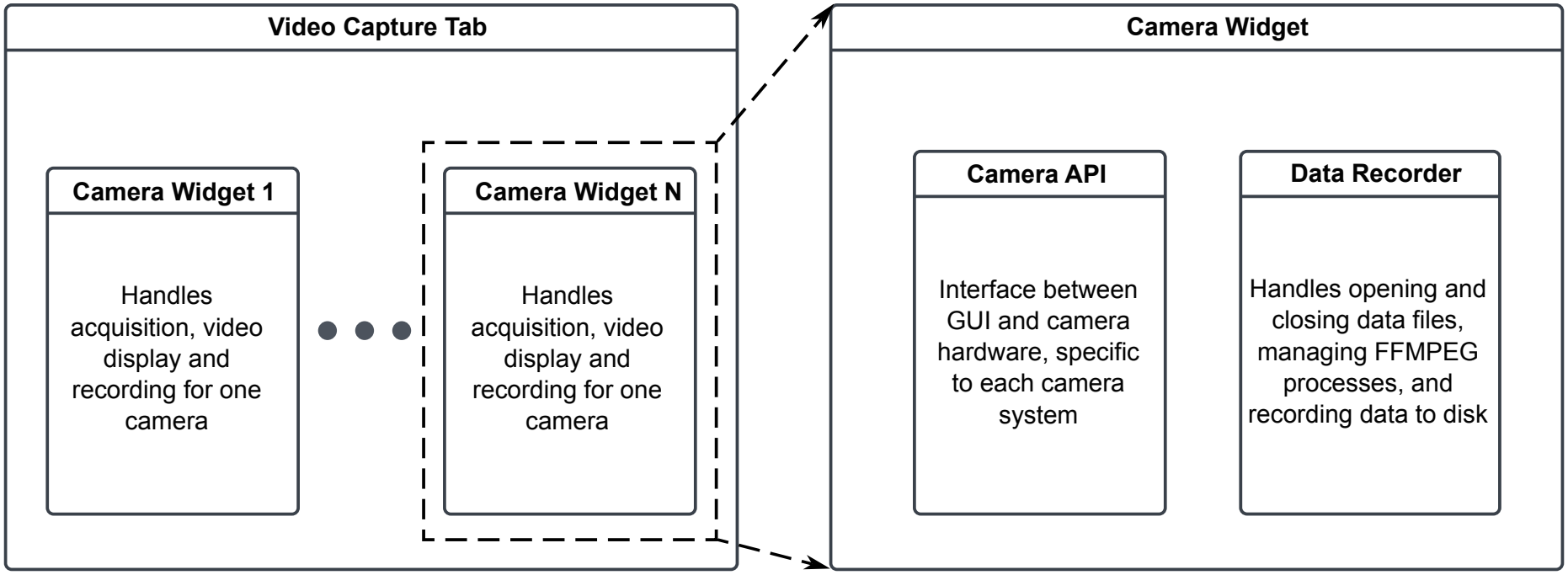
Software components. Organisation of the main software components involved in video acquisition. Each component is a separate Python class.

### Camera API

Camera APIs provide an abstraction layer between the GUI and the hardware-specific Python modules needed to interact with each manufacturer’s cameras (e.g. the PySpin package used to control FLIR cameras). This allows the GUI to work with different camera systems, while providing a well-defined interface between the GUI and camera hardware which simplifies implementing support for new cameras. Each camera system supported by pyMultiVideo has its own camera API module, with camera APIs currently implemented for FLIR and Ximea cameras.

Adding support for a new camera system requires creating a corresponding camera API module. The technical reference for camera APIs is defined by a generic_camera module, which contains a *GenericCamera* class that specifies the methods required for interaction with the GUI, as well as functions to get the serial number of connected cameras, and instantiate a camera object given a serial number. See the [camera API](https://pymultivideo.readthedocs.io/en/latest/user-guide/camera-api/) documentation for more information.

### Parallel Processing

pyMultiVideo’s architecture aims to balance simplicity of the code-base with performance. For simplicity the GUI is single threaded, with parallel processing used only for compressing and saving video data (Figure 4). Python’s *subprocess* module is used to spawn a separate process for each camera, which runs an FFMPEG instance used to compress and save the camera’s video feed. FFMPEG is configured to use the GPU for video compression if available, dramatically reducing CPU load. To avoid the main GUI thread pausing while it waits for the FFMPEG process to accept the new frames, a Threadpool asynchronously executes the function calls that pipe newly acquired frames to FFMPEG.

**Figure 4:**
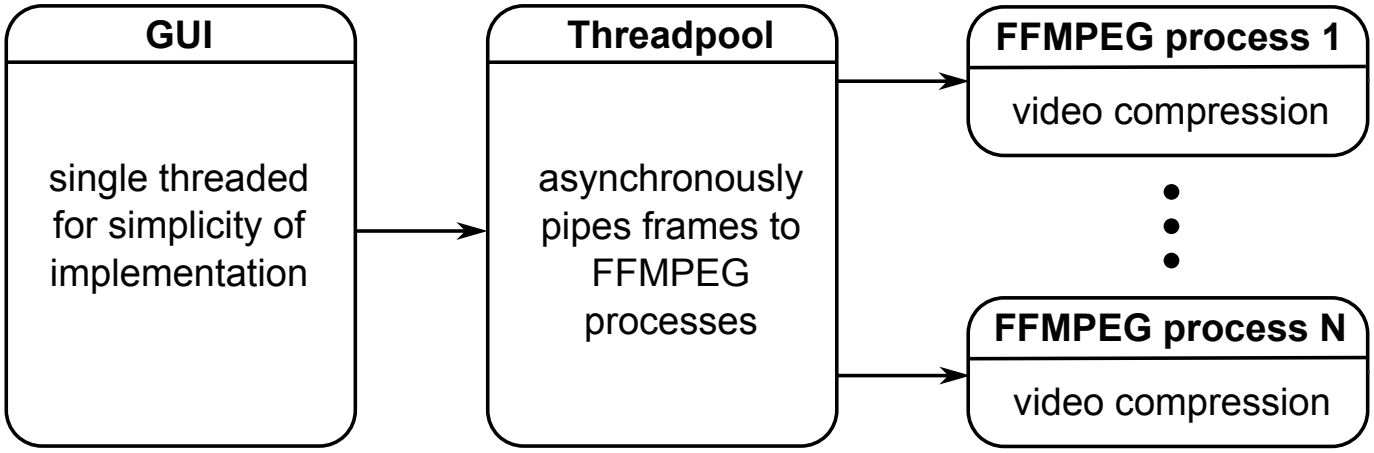
Parallel processing. Diagram illustrating the use of parallel processing in pyMultiVideo.

Another aspect of the software which is important for performance is the use of camera frame buffers implemented by the camera system’s control software (e.g. PySpin/Spinnaker for FLIR cameras), which temporarily store frames sent by the camera until they are retrieved by Python. This allows the GUI’s refresh rate - which determines how often it fetches new images from the camera, to be lower than the camera frame rate without frames being dropped.

The software is designed to ensure that it does not fail silently if it is unable to process incoming video fast enough to avoid dropping frames. Dropped frames can occur for two reasons: First, if the GUI refresh rate drops to a point where the camera frame buffers overflow between Python fetching new images, or second, if FFMPEG cannot compress the videos as fast as they are being generated. If dropped frames occur for either reason, a warning is displayed in the video overlay indicating the cause of the dropped frames. Additionally, the number of dropped frames is stored in the video’s metadata file, and the places in the video where they occurred can be identified from the frame numbers in the frame info file.

### Performance Evaluation

To quantify software performance we systematically varied a range of acquisition and encoding parameters and evaluated the fraction of dropped frames in recorded videos. Figure 5A shows performance as a function of number of cameras and acquisition frames per second (FPS), using the default video encoding parameters (h264 compression, *fast* encoding speed preset, no downsampling). With six cameras recording simultaneously no frames were dropped at up to 90 FPS, with the fraction of dropped frames increasing at higher frame rates and reaching 20% at 150FPS. With 4 cameras recording simultaneously no frames were dropped up to the cameras maximum frame rate of 150FPS. We note that the computer used for these experiments was relatively old (i7-9700 CPU, RTX2060 GPU, released 2019) and higher performance would be anticipated on a more powerful machine.

**Figure 5:**
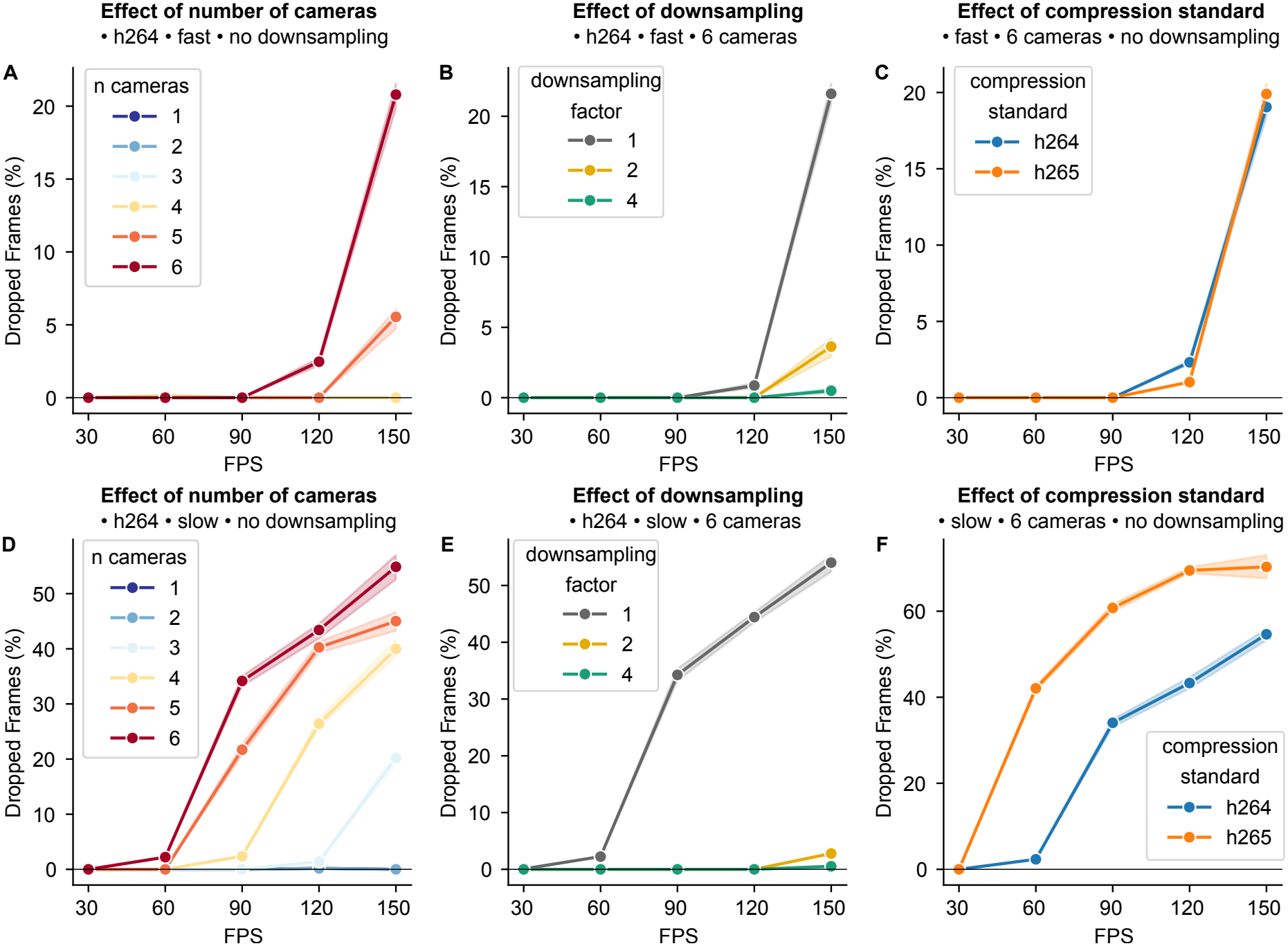
Performance quantification. Evaluation of pyMultiVideo performance, quantified as the percentage of dropped frames, as a function of different acquisition parameters. Parameters varied across experiments were: Frames per seconds (FPS), number of cameras, encoding speed (*fast* or *slow*), spatial downsampling factor (1, 2, or 4), compression standard (*h264* or *h265*). Panels **A** - **C** show effect of number of cameras, downsampling and compression standard when encoding speed was set to *fast*. Panels **D** - **F** show effect of same variables when encoding speed was set to *slow*.

To quantify how FFMPEG encoding parameters affected performance, we systematically varied spatial downsampling, the compression standard (*h264* vs *h265*) and encoding speed presets (*fast* vs *slow*). Encoding speed had a large effect on performance, with frames dropped above 30FPS with six cameras when slow encoding speed was used (Figure 5D). Slower encoding speed can achieve higher video quality for a given file size, but does so at the cost of greatly increased compute demands. Compression standard had little effect on performance when the encoding speed was fast (Figure 5C), but a substantial effect on performance when the encoding speed was slow (Figure 5F), where *h264* compression resulted in less dropped frames compared with *h265*. This is likely because at fast encoding speed the bottleneck causing dropped frames is fetching images from the cameras, whereas at slow encoding speed the bottleneck is video compression. Applying spatial downsampling to the videos (implemented by FFMPEG prior to compression), could dramatically reduce the rate of dropped frames (Figure 5B,E), though at the cost of reducing video resolution. Qualitatively similar results were obtained on a second computer with i7-6700 CPU and RTX A400 GPU (Figure S1).

## Discussion

We developed an open source software tool for acquiring, synchronising and compressing video from multiple cameras for scientific applications. The principal strengths of the software are; a user friendly GUI designed for high-throughput multi-camera experiments, support for multiple synchronisation methods, online video compression at high frame rates using the GPU, detailed online documentation, and a Python code base structured to facilitate adding support for new camera systems.

A variety of alternative software solutions exist for video acquisition. Scientific cameras typically come with viewer software with basic recording functionality, e.g. Spinview for FLIR or CamTool for Ximea cameras. These are primarily designed for testing and configuring cameras and often lack features useful for experimental applications such as support for multi-camera recordings, GPU-based video encoding, or saving GPIO pin-states and video metadata. Many commercial software packages exist for recording from multiple CCTV cameras, however these cameras typically have relatively low frame rates, and do not support external frame triggering or GPIO input needed for accurate synchronisation. Commercial software for multi-camera acquisition using scientific camera systems does exist, e.g. [StreamPix](https://www.norpix.com/products/streampix/streampix.php), however as specialist commercial software it requires an expensive licence.

Several open source tools have been developed that support scientific video acquisition. Bonsai ^14^ is a visual programming language for creating data processing workflows which can be used for video acquisition. Indeed, prior to developing pyMultiVideo we used a workflow combining Bonsai and FFMPEG for video acquisition ([Github](https://github.com/ThomasAkam/Point_Grey_Bonsai_multi_camera_acquisition)). In our experience, this Bonsai-based video acquisition pipeline was less user-friendly for configuring and running experiments than pyMultiVideo. This may in part reflect our limited Bonsai programming skills, but also reflects inherent trade-offs between flexibility and simplicity. As a powerful general purpose tool for scientific data processing and experiment control, Bonsai supports a wide range of complex use cases, while pyMultiVideo is a single purpose tool for video recording, permitting a streamlined GUI design optimised for this application.

Dedicated open source software packages for scientific video acquisition have also been developed ^7–11^, with varying user interfaces, features sets and documentation (Table 1). Our motivation for developing pyMultiVideo was that we required a set of features not found together in an existing package, specifically; a user-friendly GUI for efficiently configuring and running high-throughput experiments, the ability to read GPIO pin-states for synchronisation, and online GPU-based h264 video compression. To maximise the likelihood of the software being useful for other groups, we have structured and documented the code to facilitate adding support for new camera systems, and implemented detailed online user documentation.

**Table 1:**
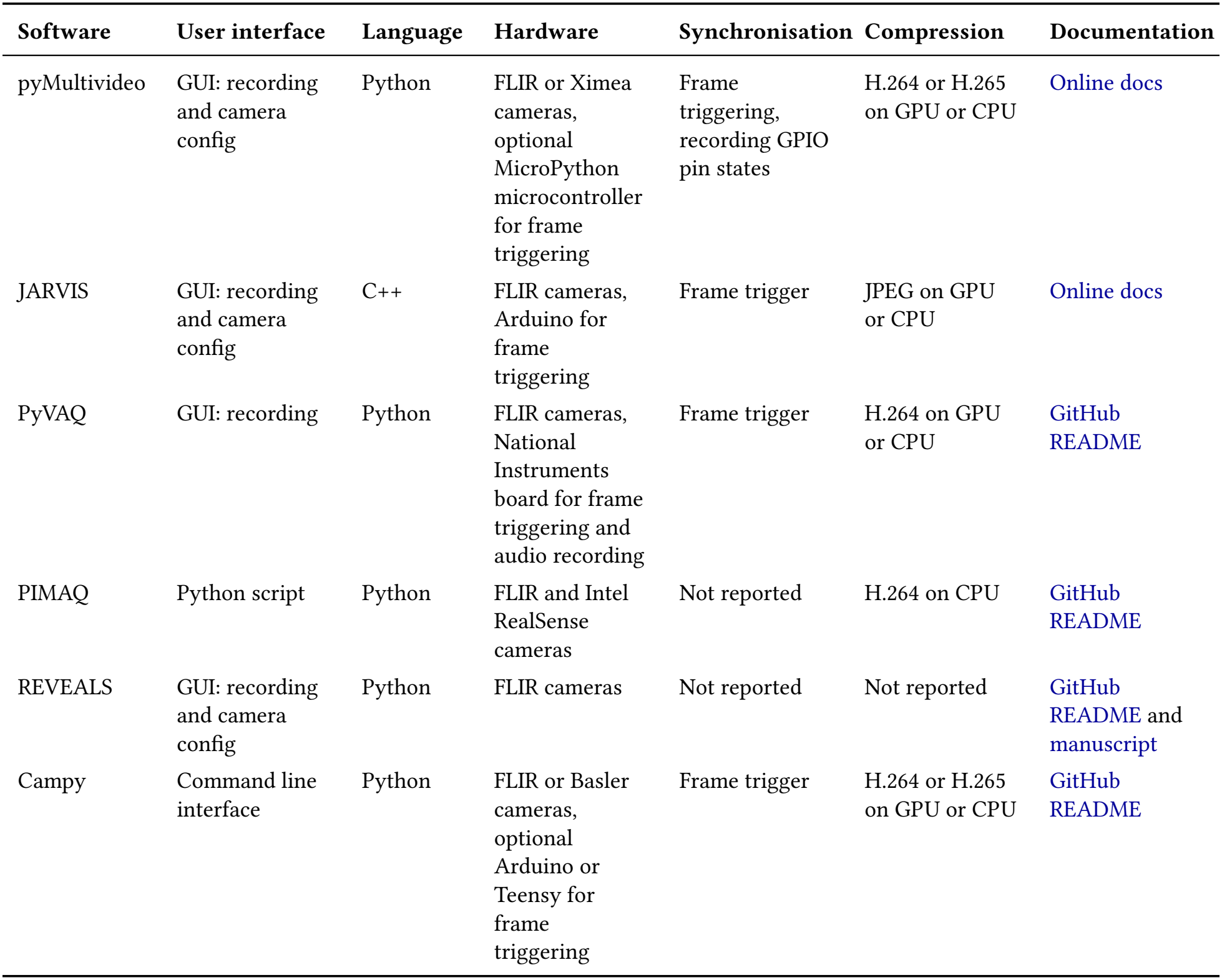
Comparison of software packages for video acquisition.

| **Software** | **User interface** | **Language** | **Hardware** | **Synchronisation** | **Compression** | **Documentation** |
|---|---|---|---|---|---|---|
| pyMultivideo | GUI: recording and camera config | Python | FLIR or Ximea cameras, optional MicroPython microcontroller for frame triggering | Frame triggering, recording GPIO pin states | H.264 or H.265 on GPU or CPU | Online docs |
| JARVIS | GUI: recording and camera config | C++ | FLIR cameras, Arduino for frame triggering | Frame trigger | JPEG on GPU or CPU | Online docs |
| PyVAQ | GUI: recording | Python | FLIR cameras, National Instruments board for frame triggering and audio recording | Frame trigger | H.264 on GPU or CPU | GitHub README |
| PIMAQ | Python script | Python | FLIR and Intel RealSense cameras | Not reported | H.264 on CPU | GitHub README |
| REVEALS | GUI: recording and camera config | Python | FLIR cameras | Not reported | Not reported | GitHub README and manuscript |
| Campy | Command line interface | Python | FLIR or Basler cameras, optional Arduino or Teensy for frame triggering | Frame trigger | H.264 or H.265 on GPU or CPU | GitHub README |

Our software has several limitations. Although we provide detailed step-by-step installation instructions, installation is still somewhat cumbersome as it requires creating a dedicated Python environment and downloading and installing packages from camera manufacturer websites. Though key camera settings can be configured from the GUI (fps, gain, exposure and trigger mode), configuration of other camera settings is not currently supported. While the software achieves high performance in terms of the total frame-rates recorded without dropping frames, the GUI can become laggy in high load conditions. py-MultiVideo currently only supports cameras from FLIR and Ximea, and has been extensively tested only with FLIR Chameleon 3 cameras, though some testing has used FLIR Blackfly and Ximea xiQ cameras. As pyMultiVideo is open source, users can modify the software to address these limitations, and we would particularly welcome development of APIs to extend support to a wider range of camera hardware.

In summary, pyMultiVideo is a useful and versatile tool for scientific video acquisition, which we hope will facilitate the ever expanding use of video data in understanding behaviour.

## Methods

### Resources

- Online documentation: https://pymultivideo.readthedocs.io
- User discussion board: https://github.com/orgs/pyMultiVideo/discussions
- Software code repository: https://github.com/pyMultiVideo/code
- Documentation repository: https://github.com/pyMultiVideo/docs

### Performance evaluation

We evaluated application performance by quantifying the fraction of dropped frames during recording as a function of recording parameters including the number of cameras, frame rate, video compression standard, encoding speed, and spatial downsampling. Experiments were run on two different computers. Computer 1 (Figure 5) had an Intel i7-9700 CPU, NVIDIA RTX2060 GPU, 32GB of RAM. Computer 2 (Figure S1) had an Intel i7-6700 CPU, NVIDIA RTXA400 GPU, 32GB of RAM. All performance evaluations used Chamleon3 CM3-U3-13Y3C-CS cameras connected to the computer by USB3.

Parameters were systematically varied across recordings using a Python script which automatically launched the GUI from the command line, configured the recording parameters, and recorded video for a duration of 1 minute per condition. Numbers of recorded and dropped frames were extracted from the recording metadata files. The test script and other code needed to configure, run and plot performance evaluations are contained in the [test](https://github.com/pyMultiVideo/code/tree/main/test) folder in the software code repository.

## Acknowledgements

Work was supported by Wellcome Trust Career Development Award 225926/Z/22/Z to TA. For the purpose of open access, the author has applied a CC BY public copyright licence to any Author Accepted Manuscript version arising from this submission.

## Author contributions

A.A. and T.A. designed and implemented the software, analysed data and wrote and edited the manuscript.

## Data Availability

The data and code used to generate the performance evaluation figures are available at: https://github.com/pyMultiVideo/manuscript

## Supplementary Figures

**Figure S1:**
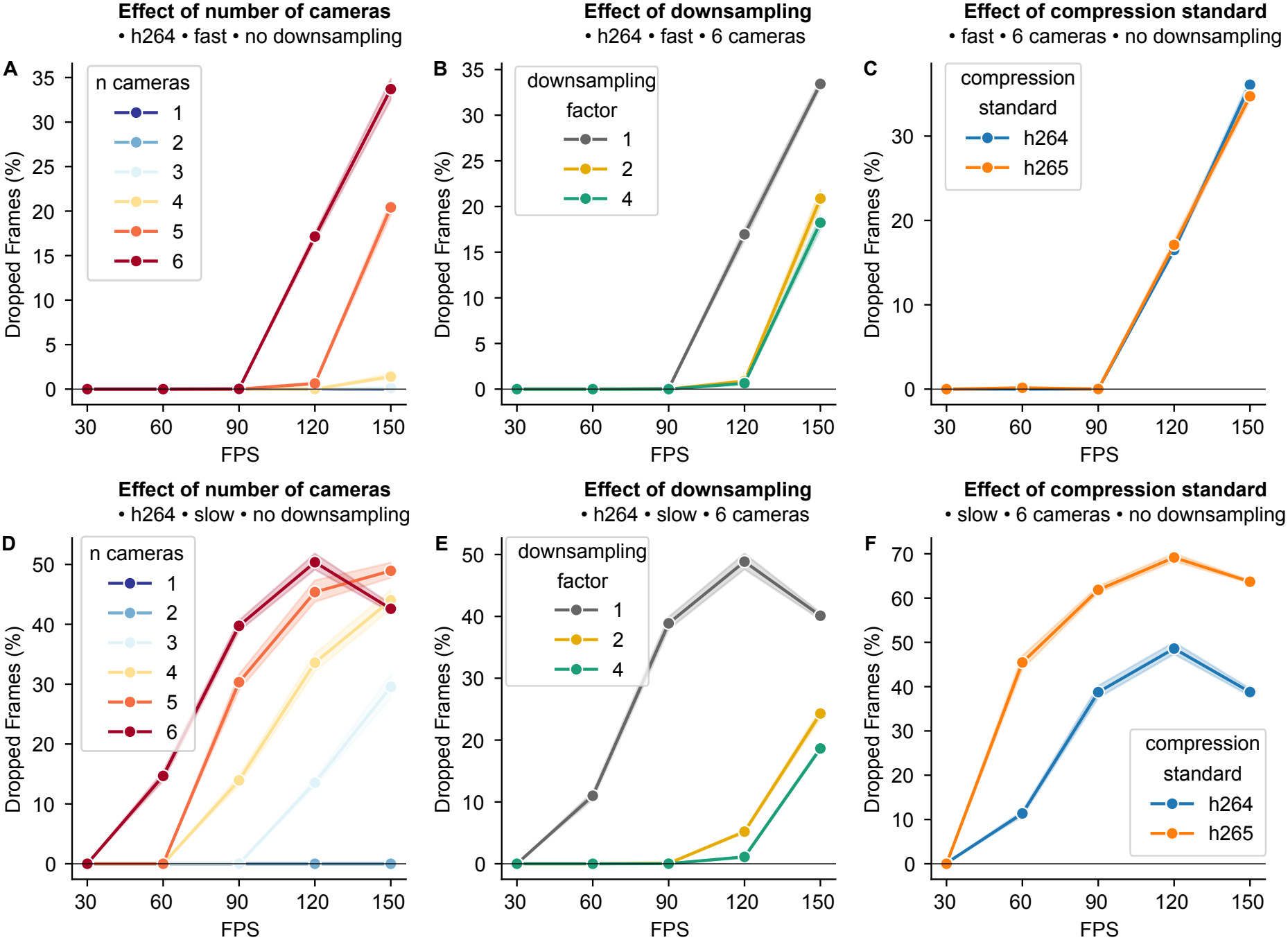
Performance quantification on second computer. Evaluation of pyMultiVideo performance on a second computer (i7-6700 CPU, RTX A400 GPU, 32GB RAM). Performance was quantified as the percentage of dropped frames, as a function of different acquisition parameters. Parameters varied across experiments were: Frames per seconds (FPS), number of cameras, encoding speed (*fast* or *slow*), spatial downsampling factor (1, 2, or 4), compression standard (*h264* or *h265*). Panels **A** - **C** show effect of number of cameras, downsampling and compression standard when encoding speed was set to *fast*. Panels **D** - **F** show effect of same variables when encoding speed was set to *slow*.

